# MIMEco: Multi-objective metabolic modeling to predict and explain pairwise interactions

**DOI:** 10.1101/2025.11.25.690486

**Authors:** Anna Lambert, Samuel Chaffron, Damien Eveillard

## Abstract

Genome-scale metabolic models (GEMs) provide a mechanistic framework for quantitatively exploring an organism’s physiology and its interactions with the environment. However, their use in microbial community modeling has been limited since it usually relies on community-wide objective functions or the integration of community abundance data. Nevertheless, predicting microbial interactions is crucial for understanding ecosystem dynamics, and thus requires dedicated methods. Here, we introduce MIMEco (Metabolic Interaction Modeling in Ecosystems), an open-source Python package that predicts pairwise microbial interactions through multi-objective optimization. MIMEco does not require abundance data, provides a user-friendly workflow, and predicts interaction types, strengths, and exchanged metabolites directly from Pareto fronts. In this study, MIMEco is validated by replicating experimental results of an *in vitro* co-culture of two auxotrophic *Escherichia coli* strains. By combining mechanistic inference with usability, MIMEco makes metabolic interaction modeling accessible to non-specialists while remaining flexible for experts, establishing a versatile platform for advancing metabolic ecology across biomedical and environmental fields.

**IMPORTANCE:** Deciphering microbial interactions through *in vitro* experiments is often costly and limited in scope, while existing bioinformatics tools can be complex to use or limited by actual sample data. MIMEco offers an open-source and user-friendly multi-objective framework for predicting and quantifying pairwise microbial ecosystem interactions and nutrient exchanges. We demonstrate its reliability by replicating a well-characterized laboratory experiment, showing that MIMEco can accurately reflects real biological behavior. This tool improves the community’s ability to explore and understand metabolic interaction mechanisms underpinning microbial ecosystems.

## INTRODUCTION

Progress in microbial ecology are essential across research domains, such as health [1], agriculture [2], and climate change [3]. High-throughput sequencing has revolutionized the field, enabling the rapid and affordable profiling of microbial communities in diverse environments [4, 5]. While these developments have revealed central links between microbial taxa and specific conditions [6, 7], the inference and understanding of causal mechanisms still mainly relies on expensive and laborious *in vitro* experiments.

To address this need, kinetic models have been developed to simulate dynamic microbial interactions and predict *in silico* cellular behavior. Despite their potential, these models are complex to build and require significant experimental data and computational resources [8]. In contrast, genome-scale metabolic models (GEMs), which assume quasi-stationary conditions, offer a more scalable and accessible framework for exploring microbial metabolism and ecological interactions [9]. This modeling approach has demonstrated accurate predictions when simulating single-organism GEMs, where the standard goal is to maximize a growth-rate objective [10], as higher growth rates are often linked to increased fitness [11]. When applied to microbial community models, this framework was initially extended by maximizing total community biomass production [12], providing a computationally efficient method. However, this often results in unrealistic dominance of fastest-growing copiotrophic species, overlooking the coexistence of slower-growing oligotrophic organisms that are often observed in nature.

To address this limitation, several strategies incorporate species abundance data into the community objective [13]. For example, the Microbiome Modeling Toolbox (MMT) [14, 15] enables maximizing abundance-weighted biomass production while preserving the species ratios observed in real ecosystems. However, relative abundance does not directly reflect growth rate, and using it as a community objective weight can artificially limit the growth of lower abundant taxa to support the growth of higher abundant taxa [13]. MICOM [16] improves on this by using relative abundances and L2 regularization to estimate growth rates consistent with biological data, as long as the objective value of the community is suboptimal. While these approaches improve observation fitting, they rely on empirical measurements and are constrained to sampled community composition, limiting their predictive power under various conditions. Other methods aim to identify potential metabolic interactions without (relative) abundance data. For example, SMETANA [17] predicts metabolic dependencies by evaluating whether a species can grow depending on the available community-derived metabolites. However, for pairwise interaction prediction, it does not simulate simultaneous community growth, and predicted exchanges are thus not derived from realistic community-level metabolic behavior. Additionally, it lacks the ability to integrate detailed, quantitative medium compositions, which can critically influence metabolic interactions in ecosystems [18].

More mechanistic, non-parametric approaches, such as OptCom [19], utilize bi-level optimization to balance individual growth objectives with a global ecosystem objective. While this enables more realistic modeling of community behavior, it is computationally intensive and not accessible as a ready-to-use tool. The pairwise modeling framework from the MMT [14, 15] provides a practical alternative by combining two organisms into a shared model and manually inferring trade-offs between their objectives via sequential biomass constraints. This results in a Pareto front that can be interpreted to predict the type of interaction between two organisms (e.g., mutualism, competition). However, because it does not support formal multi-objective optimization, it cannot be readily scaled to larger ecosystems.

To fill the gaps of above approaches (see Table **1** for a concise overview of methods), we present MIMEco (Metabolic Interaction Modeling in Ecosystems), a user-friendly and open-source Python package (https://github.com/Anna-cell/mimeco) for predicting pairwise microbial interactions from GEMs. This package implements the framework presented in Lambert et al. [20], as it employs the use of formal multi-objective optimization to quantify trade-offs between species growth, infer interaction types (e.g., competition, neutrality, and diverse forms of mutualism), compute interaction scores, and identify exchanged metabolites, all without requiring abundance data. The tool is designed to be accessible and well-documented (https://mimeco.readthedocs.io/en/latest) to facilitate multi-objective modeling for non-experts.

**TABLE 1.** Comparison of metabolic modeling tools for microbial communities.

| Criterion | MMT - Microbial community modeling | MICOM | Smetana | OptCom | MMT - Pairwise modeling | MIMEco |
| --- | --- | --- | --- | --- | --- | --- |
| **Modeling assumption, from more to less parametric** | Community biomass optimization (abundance-weighted) | Cooperative tradeoff (abundance-weighted and regularized) | Individual growth tested with minimal interspecies support | Bi-level optimization (community + individual) | Manual trade-off via sequential biomass constraints | Multi-objective |
| **Independent of biological sample** | No (requires sample-specific abundance data) | No (requires sample-specific species abundance) | Yes | Yes | Yes | Yes |
| **Community size** | Community-wide | Community-wide | 35-40 organisms [21] | 4 [19] | Pairwise | Pairwise, but perspective to scale to a dozen [20] |
| **Medium personalization** | Quantitative | Quantitative | Qualitative | Quantitative | Quantitative | Quantitative |
| **User-friendly (standalone tool with documentation)** | Yes | Yes | Yes | No (optimization framework, requires manual implementation) | Yes | Yes |
| **Software accessibility** | Limited (Open source code, but requires MATLAB) | Open source | Open source | No official package | Limited (Open source code, but requires MATLAB) | Open source |

For validation and performance estimation, MIMEco predictions were confronted with an *in vitro* experiment considering two auxotrophic *Escherichia coli* strains. MIMEco accurately reproduced their growth impairment and recovery in mono- and co-culture, predicted the expected positive interaction score and type, as well as the cross-feeding of missing amino acids.

## RESULTS

### Multi-objective metabolic modeling to qualify and quantify pairwise interaction

Conventional metabolic modeling typically focuses on optimizing a single objective function, such as biomass production [22], and inferring an optimal solution for a given set of conditions. In contrast, multi-objective optimization allows simultaneous optimization of multiple objectives, revealing trade-offs among potentially competing metabolic goals. The outcome of such an analysis is called a Pareto front, which represents the set of solutions in which no objective can be improved without compromising another objective [23]. When applied to communities, this enables the exploration of ecologically realistic metabolic configurations that account for the performance of all organisms. MIMEco leverages this optimization principle, originally developed in Operational Research, to qualify and quantify interactions between two organisms. Given two input genome-scale metabolic models (GEMs) and a defined metabolic environment, MIMEco constructs a compartmentalized ecosystem model in which both organisms share a common extracellular pool. This shared compartment enables the exchange of metabolites. The influx of metabolites into the shared pool is constrained by the user-defined medium, ensuring control over environmental inputs.

To assess the impact of the interaction on an organism fitness, MIMEco first simulates each organism metabolic behavior in monoculture in a given medium to determine its maximum objective value (e.g., growth rate). These values are then used to normalize the axes of the Pareto front, setting “1” as the reference for performance in isolation. This step, illustrated in Figure **1**A, forms the basis for subsequent inference of interaction type, interaction score, and cross-fed metabolites.

**FIG 1.**
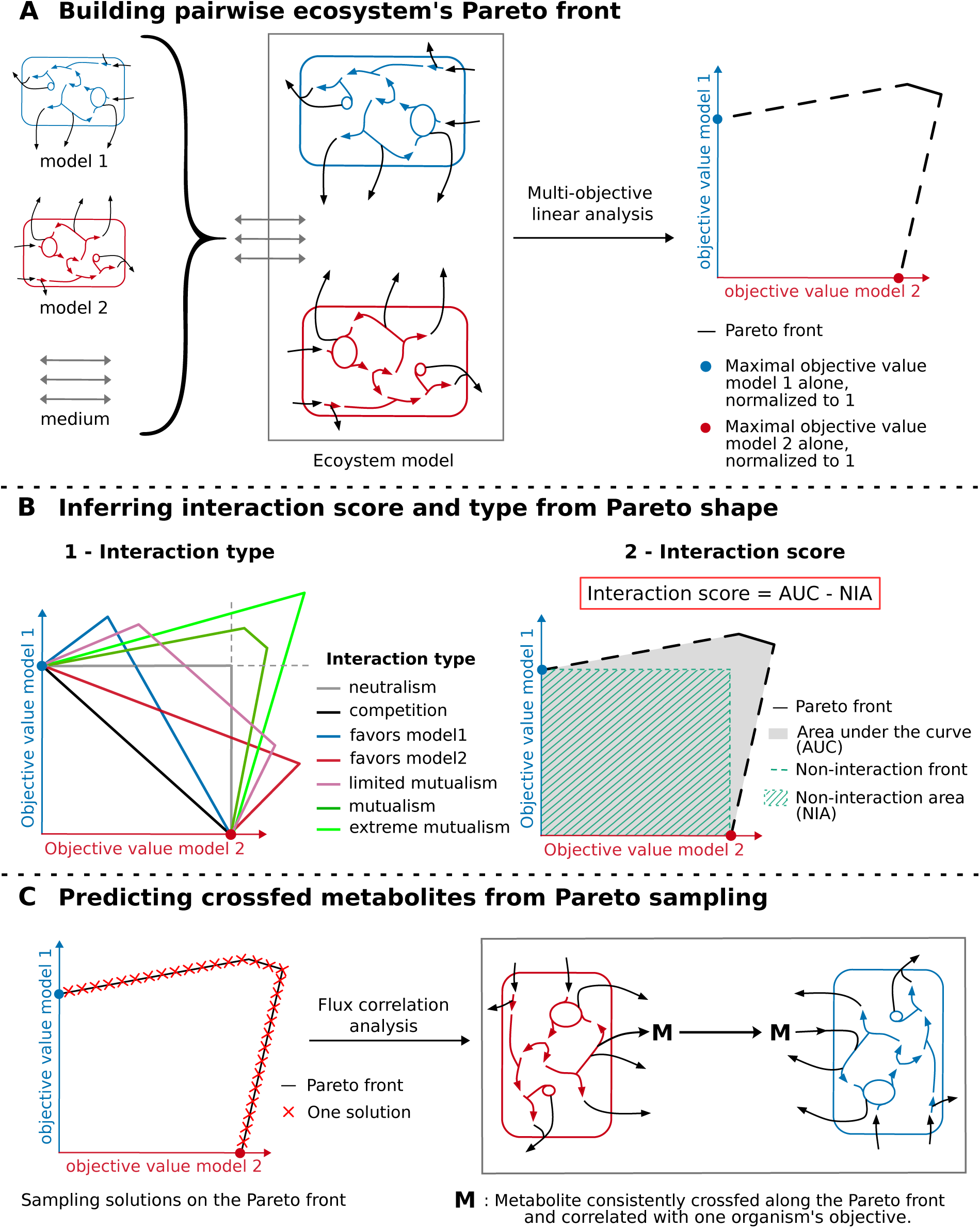
Schematic rationale of MIMEco’s automated analysis. **A)** Starting from user-provided input (two GEMs and a defined medium), MIMEco constructs a pairwise ecosystem model to simulate metabolic interactions in a controlled environment. It computes the maximal objective value for each GEM in monoculture and performs a multi-objective optimization, yielding a normalized Pareto front. **B)** From this Pareto front, the **interaction_score_and_type()** function infers **1.** the interaction type, based on the shape of the Pareto front (see Methods) and **2.** an interaction score, computed from the difference between the Pareto front’s AUC and the non-Interaction front area. **C)** The **crossfed_metabolites()** function samples solutions along the Pareto front to identify metabolites consistently exchanged between models and whose exchanges are correlated with variations in their objective values.

### Interaction type

Because each organism’s optimal objective value in monoculture is normalized to 1, deviations from this value in the community context provide insight into the nature of the interaction. Suppose an organism achieves a higher objective value within the community than it does alone. In that case, this indicates that it benefits from the presence of the other species, suggesting a commensal or mutualistic interaction. Conversely, a reduction in objective value points to competition. When two organisms yield a single Pareto-optimal solution in which both objective values match their respective monoculture optima, the interaction is classified as neutralism, indicating that neither organism significantly influences the metabolic performance of the other [20].

Figure 1 B1 shows representative Pareto fronts for seven interaction types (neutralism, competition, favors model1, favors model2, limited mutualism, mutualism, extreme mutualism), each based on trade-offs observed in shared-resource environments. The detailed description of how these interaction types are obtained is provided in Methods, Table **4**.

### Interaction score

While interaction types offer a qualitative classification of ecological relationships, the interaction score provides a complementary quantitative overview of the interaction. To define this score, we use the neutral non-interaction front (i.e., the neutrality interaction type), where both organisms reach but are limited to their monoculture optima. The area beneath it defines the Non-Interaction Area (NIA). MIMEco calculates the interaction score as the difference between the Area Under the Curve of the Pareto front (AUC) and the NIA, as shown in Figure 1 B2. A positive score indicates that the ecosystem allows one or both organisms to exceed their monoculture objective values, as in commensal or mutualistic interactions. Conversely, a negative score indicates decreased metabolic performance, often associated with competitive interactions [20]. The magnitude of the score reflects the overall metabolic benefit or cost of the interaction relative to independent growth.

It is important to recognize that positive and negative interaction scores are not directly comparable in magnitude, as the minimum score is limited. Specifically, because of the convexity of the Pareto front, the worst-case competitive interaction happens when the front includes only the two monoculture optima (i.e., the points (1, 0) and (0, 1)), forming a straight diagonal line (shown as the example competition interaction type in Figure 1 B1). In this scenario, the area under the curve (AUC) is 0.5, and the non-interaction area (NIA) remains 1. Consequently, the lowest possible score is −0.5. Conversely, positive scores are unbounded, as mutualistic interactions can allow organisms to exceed their monoculture performance by a significant margin. Therefore, negative and positive scores should be considered within their respective ranges and not as symmetric measures of interaction strength. Both interaction type and interaction score are returned by the MIMEco function **interaction_score_and_type()**

### Sampling solutions on the Pareto front to identify consistently cross-fed metabolites

A key advantage of GEMs is their ability to uncover potential mechanisms underpinning interactions [24, 25]. To harness this potential, we developed a method to identify metabolic exchanges that correlate with changes in an organism’s objective value, whether those changes indicate improvements or impairments in performance.

This analysis starts with sampling 10,000 solutions along the Pareto front (as shown in Figure 1C), each representing a unique feasible community-level metabolic behavior. For each reaction, including exchange and objective reactions, pairwise flux correlations are calculated. We then identify candidate cross-fed metabolites (Figure 1 C) as those that meet two conditions: (i) they are consistently exchanged (i.e., secreted by one organism and taken up by the other) across a significant portion of the sampled solutions, and (ii) their exchange flux is correlated (positively or negatively) with the objective value of at least one of the organisms. This approach allows the prediction of functionally relevant metabolite exchanges that likely drive the interaction dynamics.

This analysis is implemented in the MIMEco function **crossfed_metabolites()**, which returns a dataframe summarizing how each inferred cross-fed metabolite meets the criteria defined above. Additionally, the analysis conditions can be adjusted via user-defined threshold parameters, allowing flexible control over sample size, correlation strength, and exchange consistency requirements (see the cross-fed metabolites section in Methods). A plotting option enables the visualization of the evolution of transport fluxes along the Pareto front during sampling for the inferred cross-fed metabolites, facilitating understanding and interpretation of the predictions. An illustration of these plots and an example analysis are available in the supplementary materials.

### Specialized functions for the human small intestinal environment

The method used in MIMEco builds on earlier research on metabolic interactions between the small intestinal microbiome and human enterocytes [20]. In this context, the small intestinal epithelial cell model developed by Sahoo et al. [26] has been extensively studied through multi-objective metabolic modeling. However, because of its unique structure, which includes two external compartments and a specific metabolite namespace, it is not directly compatible with the standard MIMEco workflow.

To address this, MIMEco includes two dedicated functions specifically designed to analyze interactions between a gut microbe model and the pre-integrated adapted small intestinal enterocyte model: **enterocyte_interaction_score_and_type()** and **enterocyte_crossfed_metabolites()**. These functions mirror the behavior and outputs of their standard MIMEco counterparts but require only a single microbial GEM as input, with the enterocyte model embedded in the analysis pipeline.

### Validation of MIMEco predictions using *in vitro* experiments

Previous work has shown that known probiotic strains tend to achieve higher interaction scores than non-probiotic strains when modeled in ecosystems containing small intestinal epithelial cells [20], providing initial support for the biological relevance of MIMEco predictions. While these results are promising, further validation is needed to assess the method’s reliability in predicting and explaining metabolic interactions.

To this end, we reproduced the experimental setup described by Jo et al. [27], which introduces the BioMe device: a co-culture platform comprising two compartments separated by a semipermeable membrane. This design allows for diffusion-based metabolic exchange between organisms while enabling independent measurement of each strain’s growth rate. The system provides an ideal framework for evaluating MIMEco’s ability to predict pairwise metabolic interactions in a controlled, experimentally tractable context.

In this paper, we reproduced *in silico* a co-culture experiment involving two auxotrophic derivatives of *Escherichia coli* MG1655 and compared the results with those of the *in vitro* experiment.

### In vitro co-culture of amino acid auxotrophic E. coli strains

In the protocol by Mee et al. [28], two *E. coli* MG1655 derivatives were created: a lysine auxotroph (ΔLys, with a KO on *lysA*) and an isoleucine auxotroph (ΔIle, will a KO on *ilvA*) (See Figure 2 A1). As expected, neither mutant grew in minimal medium without amino acid supplementation. Monoculture growth was restored by the addition of the corresponding missing amino acid (lysine or isoleucine).

**FIG 2.**
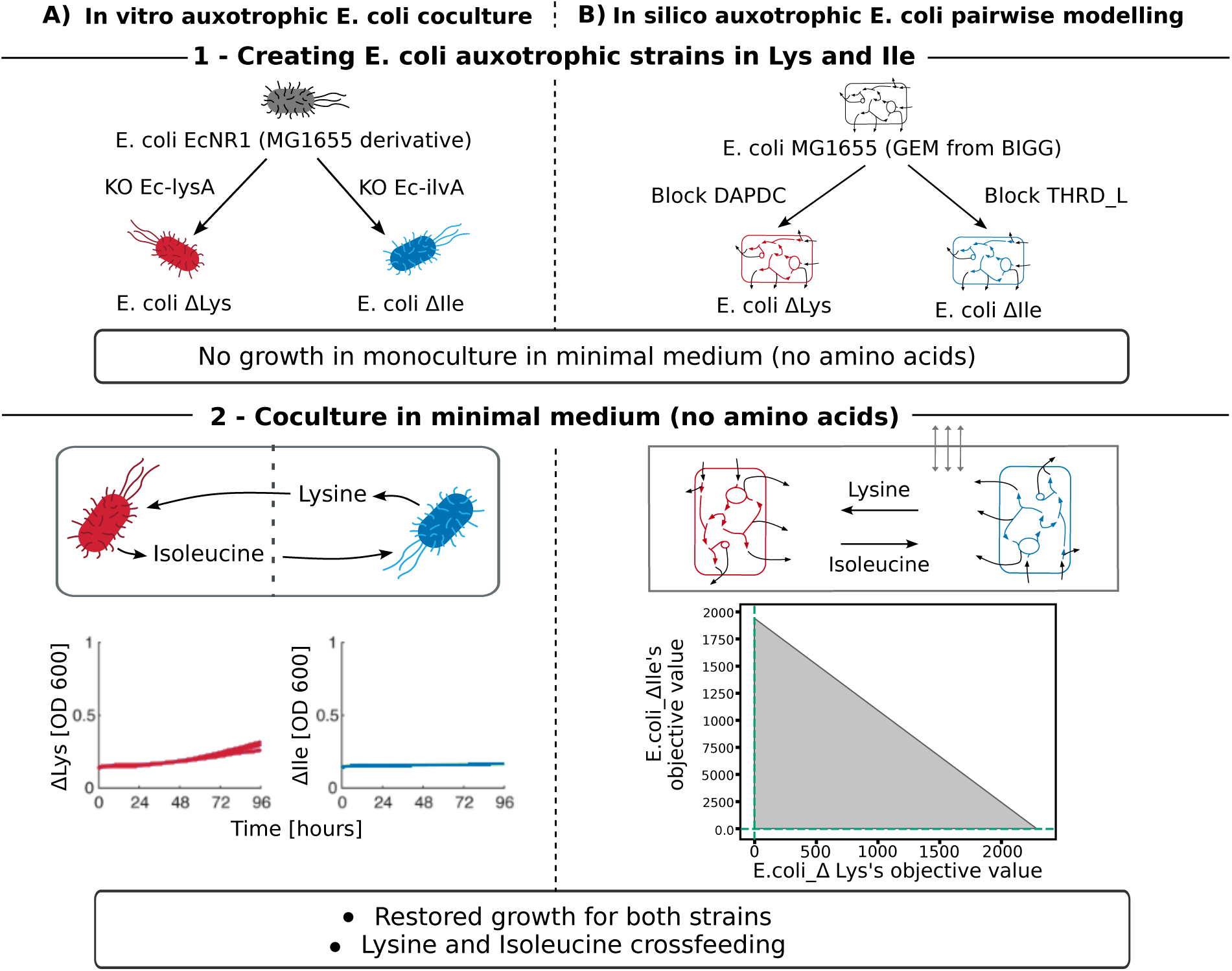
Parallel illustration of the *in vitro* auxotrophic *E. coli* co-culture experiment and its *in silico* replication. **A)** In the *in vitro* experiment: **(1)** knockout of the lysA or ilvA gene generates *E. coli* strains auxotrophic for lysine or isoleucine, respectively. These auxotrophs fail to grow in monoculture but **(2)** recover growth in co-culture via cross-feeding of the missing amino acids. **B)** The *in silico* replication: **(1)** the corresponding GEMs are modified by blocking the DAPDC or THRD_L reactions from an *E. coli* GEM to simulate lysine or isoleucine auxotrophy. **(2)** Pairwise co-culture using MIMEco restores growth and yields a strongly positive interaction score (2,283,562).

When co-cultured in a BioMe device under minimal medium conditions, ΔLys and ΔIle exhibited restored growth [27], as illustrated in Figure 2 A2. However, growth rates remained constrained by the porous membrane of the device, and the benefit of co-culture was asymmetric: ΔLys achieved a higher growth rate than ΔIle (Figure 2A). Still, ΔIle showed improved growth in co-culture relative to its monoculture without isoleucine, confirming syntrophic cross-feeding [27].

### In silico co-culture of amino acid auxotrophic E. coli strains

To reproduce this experiment *in silico*, we used MIMEco with genome-scale models of both auxotrophic strains. Starting from the high-quality BiGG model of E. coli MG1655, we blocked the DAPDC and THRD_L reactions to simulate the *lysA* and *ilvA* knockouts, respectively. Simulations were run in a minimal medium (MM), adapted from experimental conditions [28, 27].

Individual optimization of ΔLys and ΔIle in MM was infeasible, confirming the absence of viable growth solutions. Wild-type *E. coli* (WT), however, reached a maximal growth rate of 0.7109 gDW.h*^−^*^1^. To enable normalization to the monoculture optimum in MIMEco, we introduced a trace amount of the missing amino acid (0.0001 mmol.gDW*^−^*^1^.h*^−^*^1^) in a modified minimal medium (MM^+^). This allowed marginal growth in monoculture (3.4×10*^−^*^3^ gDW.h*^−^*^1^ for ΔLys and 2.9×10*^−^*^3^ gDW.h*^−^*^1^ for ΔIle), while the WT growth remained unchanged. The medium can still be considered heavily depleted in lysine and isoleucine, as their addition in moderate amounts (1 mmol.gDW*^−^*^1^.h*^−^*^1^, in the MM^++^ medium) brings each strain, WT or auxotrophic, to a maximal growth rate of 0.7388 gDW.h*^−^*^1^.

Simulating co-culture with MIMEco in MM^+^ led to restored growth for both mutants: 0.6762 gDW.h*^−^*^1^ for ΔLys and 0.677 gDW.h*^−^*^1^ for ΔIle. These values are comparable to WT growth and reflect mutualistic cross-feeding dynamics. Growth rates across all conditions are summarized in Table **2**.

**TABLE 2.** Maximal growth rates of WT and auxotrophic *E. coli* strains in different conditions. MM is the minimal medium without amino acids. MM^+^ is the minimal medium with traces of lysine and isoleucine added to enable a negligible growth of Δ Lys and ΔIle. MM^++^ is the minimal medium with moderate amounts of lysine and isoleucine supplemented.

| Model | MM | $MM^+$ | $MM^{++}$ | Co-culture $MM^+$ |
| --- | --- | --- | --- | --- |
| WT | 0.7109 | 0.7109 | 0.7388 | 0.7109<br>(Pairwise with WT) |
| ΔLys | infeasible | 0.00029 | 0.7388 | 0.6762<br>(Pairwise with ΔIle) |
| ΔIle | infeasible | 0.00034 | 0.7388 | 0.677<br>(Pairwise with ΔLys) |

This strong recovery from near-zero growth in monoculture to near-WT levels in co-culture translates into a highly mutualistic Pareto front (Figure 2 B2), validating MIMEco’€™s capacity to simulate cooperative behavior.

The inferred interaction type is “Mutualism” (See Methods). The interaction score of 2,283,562 is unambiguous concerning the nature of the interaction. Such a high score is uncommon and reflects a dramatic difference in growth potential between monoculture and co-culture, specifically in this case, where negligible monoculture growth is treated as a proxy for the absence of growth. Interestingly, the Pareto front shows a stronger positive interaction effect on the growth of ΔLys than on ΔIle, consistent with the *in vitro* observations.

Finally, MIMEco identified lysine and isoleucine as the only cross-fed metabolites. Lysine is predicted to be secreted by ΔIle and uptaken by ΔLys in 71% of the sampled Pareto solutions, with a strong correlation (0.97) between this exchange and ΔLys growth. Visualization of lysine exchange fluxes along the Pareto front (Figure S1) shows that maximal lysine exchange from ΔIle to ΔLys coincides with the peak growth rate of ΔLys. The same pattern is observed for isoleucine, which is exported by ΔLys and supports ΔIle growth (See Table **3** and Figure S1).

**TABLE 3.** Output of the crossfed_metabolites() function for the auxotrophic E. coli strains co-culture.

|  | metabolite_id | model_benefiting | proportion_exchange | proportion_model1<br>_to_model2 | proportion_model2<br>_to_model1 | correlation<br>_obj_model1 | correlation<br>_obj_model2 |
| --- | --- | --- | --- | --- | --- | --- | --- |
| 271 | ile__L | model1 | 0.71 | 0.00 | 0.71 | 0.97 | 0.35 |
| 272 | lys__L | model2 | 0.71 | 0.71 | 0.00 | 0.35 | 0.97 |

**TABLE 4.** Classification of interaction types based on ecosystem and monoculture maximal objective value.

| Interaction type | Criteria |
| --- | --- |
| Competition | Neither model achieves a higher objective value in the ecosystem than in isolation, and the interaction score is negative. |
| Neutrality | Neither model achieves a higher objective value in the ecosystem than in isolation, and the interaction score is indistinguishable from zero. |
| Favors model 1 | Model 1 achieves a higher objective value in the ecosystem than in isolation, while model 2 does not. |
| Favors model 2 | Model 2 achieves a higher objective value in the ecosystem than in isolation, while model 1 does not. |
| Limited mutualism | Both models achieve higher objective values in the ecosystem than in isolation, but not in a synchronized solution. |
| Mutualism | At least one solution exists in which both models achieve higher objective values in the ecosystem than in isolation. |
| Extreme mutualism | A single shared solution yields the maximum ecosystem objective values for both models, and these values are higher than their respective maxima in isolation. |

## Discussion

Multi-objective metabolic modeling is a powerful method for predicting microbial community interactions, but current implementations often require extensive expertise, depend on sample-specific data, or lack accessibility. With MIMEco, we offer an open-source Python package that reduces these barriers by providing a user-friendly framework to infer interaction scores, interaction types, and cross-fed metabolites. Its modular design, organized into well-documented, task-specific functions, enables users to perform standard analyses and customize the framework for their own needs, fostering community-driven development and ongoing improvement.

MIMEco enables modeling of condition-specific pairwise ecosystems by allowing users to define the medium quantitative composition and adjust the stringency of constraints. This flexibility makes it broadly applicable across biology, such as marine ecosystems, soil microbiomes in agriculture, or host-associated microbial communities in health contexts. For gut microbiota research in particular, MIMEco includes functions for simulating interactions between microbes and a small intestinal epithelial cell with exchanges to the blood compartment.

When used to replicate the co-culture *in vitro* experiment of Jo et al. [27], MIMEco produced results that closely matched those of the original study. Notably, it reproduced the enhanced growth recovery of the ΔLys strain compared to the ΔIle, validating the framework’s predictive abilities. However, it should be emphasized that this case represents an ideal scenario: *E. coli* is among the best-characterized organisms, and the examined strain benefits from a highly curated model with a well-defined metabolic network. Additionally, the replicated experiment has very simple condition change, in two almost identical organism, which highly limits the complexity of the ecosystem model. This validation aims to validate the method implemented in MIMEco when minimizing uncertainties such as model quality, namespace or reconstruction incompatibilities, or complex medium composition.

Indeed, MIMEco’s predictive power relies heavily on the quality of the input GEMs and the accuracy of medium definition. Improvements in GEM reconstruction will thus directly boost MIMEco’s performance. Currently, the tool focuses on pairwise ecosystems; however, its formal multi-objective approach has been demonstrated to scale to larger communities with 5 to 11 organisms [20], although this still requires expert handling, high-quality models, and more computational resources. Future MIMEco developments will aim to make such multi-organism ecosystem analyses more accessible and reproducible.

## Methods

### MIMEco algorithm

#### Ecosystem model

MIMEco builds ecosystem-scale models by combining two genome-scale metabolic models (GEMs) using the mocbapy package [29]. The resulting model contains a shared pool compartment that enables metabolite exchange between the two organisms.

#### Medium definition

The user defines the growth medium as a pandas Series that describes metabolite influx rates. These values set the lower bounds of the corresponding exchange reactions in the ecosystem model. Metabolites included in the input medium are referred to as *described metabolites*, while any other metabolite associated with an exchange reaction is considered *undescribed*. Because medium definition is often labor-intensive and specific to the conditions, MIMEco offers three modes for constraining exchange reactions.

1. **blocked:** Described metabolites are constrained according to the input medium. All undescribed metabolites are blocked (lower bound = 0).
2. **partially_constrained:** Described metabolites are constrained by the input medium; undescribed metabolites are assigned a default lower bound of undescribed_met_lb (default: *−*0.1).
3. **as_is:** Exchange reaction bounds are left unchanged from the original GEMs (used when no medium is provided).

#### Multi-objective optimization: Pareto inference

Multi-objective analysis is performed through mocbapy [29], using benpy, a Python adaptation of Bensolve [30, 31]. Details of the formalism are given in Lambert et al. [20]. Since MIMEco is restricted to pairwise ecosystems, the Pareto front is two-dimensional. In parallel, each organism is simulated in isolation, and its maximal objective value is saved and used to normalize its corresponding axis. These monoculture reference points are added to the Pareto front, yielding the *extended Pareto front*, which is used in all subsequent analyses.

#### Interaction type

Interaction types are classified according to the criteria summarized in Table **4**.

#### Interaction score

The interaction score *S* is defined as the difference between the area under the curve (AUC) of the normalized extended Pareto front (*AUC_P_*) and that of the corresponding non-interaction front (*AUC_NI_*):

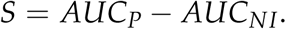

#### Crossfed metabolites

The identification of crossfed metabolites relies on a sampling of the Pareto front and a correlation analysis.

1. **Sampling of the Pareto front:** The MIMEco.utils function **pareto_sampling()** processes the Pareto front from one reference point (the maximal growth of an organism in isolation) to the other. It passes through all Pareto extreme points and samples *n* equidistant solutions, where *n* is specified by the user (default: 10,000). Each sample is obtained by constraining the ecosystem model with the two objective values from the selected Pareto solution as fixed fluxes for the objective functions, followed by computing a Flux Balance Analysis (FBA). This returns a matrix of *n* samples.
2. **Correlation:** The MIMEco.utils function **correlation()** computes a Spearman correlation on the sampling matrix, returning a correlation matrix.

A metabolite *M* is classified as *cross-fed* if it satisfies all of the following:

1. **Inverse correlation of transport fluxes.** The transport fluxes of *M* in model 1 and model 2 exhibit a correlation lower than -exchange_correlation (default: 0.5, user-defined). Negative fluxes denote uptake, positive fluxes denote secretion; thus, a negative correlation indicates potential exchange.
2. **Association with objective function.** The transport flux of *M*, in any direction and in either model, correlates with the objective function of at least one model with strength greater than biomass_correlation (default: 0.8, user-defined).
3. **Consistency of actual exchange.** The transport fluxes of *M* in model 1 and model 2 display opposite signs (one secretes, the other uptakes) in at least a proportion of lower_exchange_proportion of samples (default: 0.3, user-defined).

#### enterocyte model adaptation to MIMEco

The enterocyte model included in the MIMEco package is adapted from the small intestinal enterocyte reconstruction by Sahoo et al. (2013) [26]. Conflicts between reaction reversibility assignments and the initial reaction bounds were first resolved, with priority given to the constraints prioritized by MATLAB, to maintain the expected behavior of the original model. Incomplete gene-protein-reaction (GPR) associations were then completed. Finally, metabolite and reaction identifiers were converted to the BiGG namespace, allowing the enterocyte model to exchange metabolites with other BiGG-based reconstructions, including models generated with CarveMe. The translation script is available in the ‘paper’ folder of the MIMEco Git repository.

### Validation

#### E. coli strains

*In vitro*, the auxotroph strains of E. coli EcNR1 were obtained by knocking out key reactions of the amino acid biosynthesis pathways, as described in Mee et. al. [28]. *In silico*, the BiGG’s high-quality model of E. coli, iJO1366, was used. The biochemical reactions Ec-lysA and Ec-ilvA were respectively mapped to the model reactions DAPDC and THRD_L, whose bounds were fixed to (0, 0) in their respective auxotrophic models.

#### Nutritional environment

*In vitro*, Jo et. al. [27] realized the co-culture experiment in a minimal medium (M9 + glucose [0.4%] + thiamine [1 µg/mL] + biotin [1 µg/mL] + chloramphenicol [25 µg/mL]). *In silico*, The M9 Medium was extracted from the CarveMe database. As their medium are defined by presence (lower bound constrained at −10 for associated exchange reaction) or absence (lower bound constrained at 0 for associated exchange reaction), glucose, thiamine, biotin, and chloramphenicol were simply added as present in the original M9 medium. As lysine and isoleucine are absent from the minimal medium, but in silico traces are needed to initiate the metabolic process, the lower bound for their objective reactions was set to −0.0001.

#### MIMEco usage

In this application, MIMEco was used with the solver GUROBI.

The growth in monoculture of each E. coli strain (WT, ΔLys, ΔIle) was inferred after constraining their nutritional environment to the minimal medium:

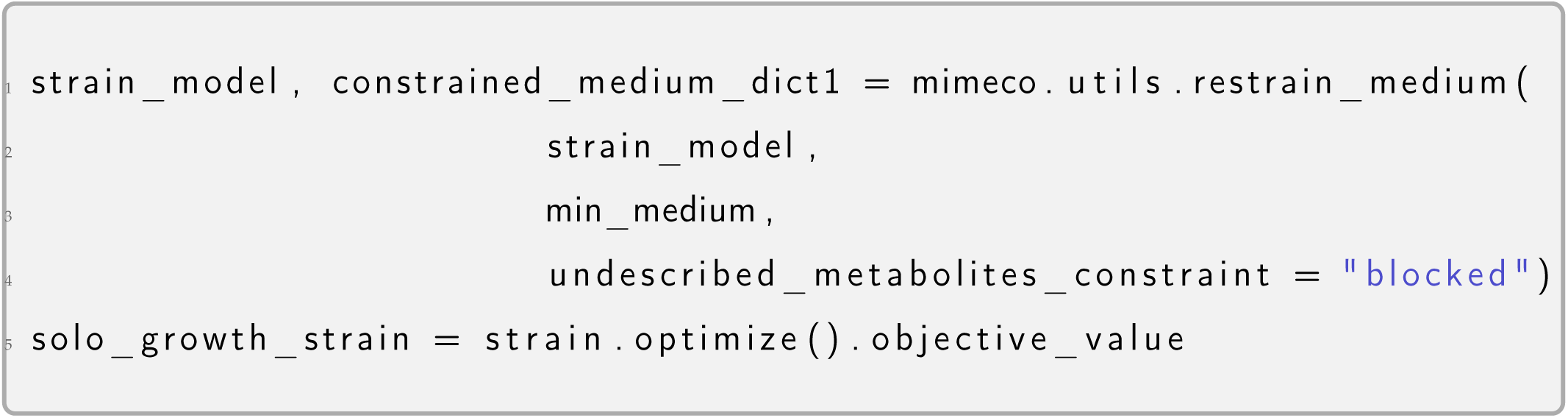

The Pareto front of the cocultured auxotrophic strain, and the interaction analysis inferred from it was obtained using:

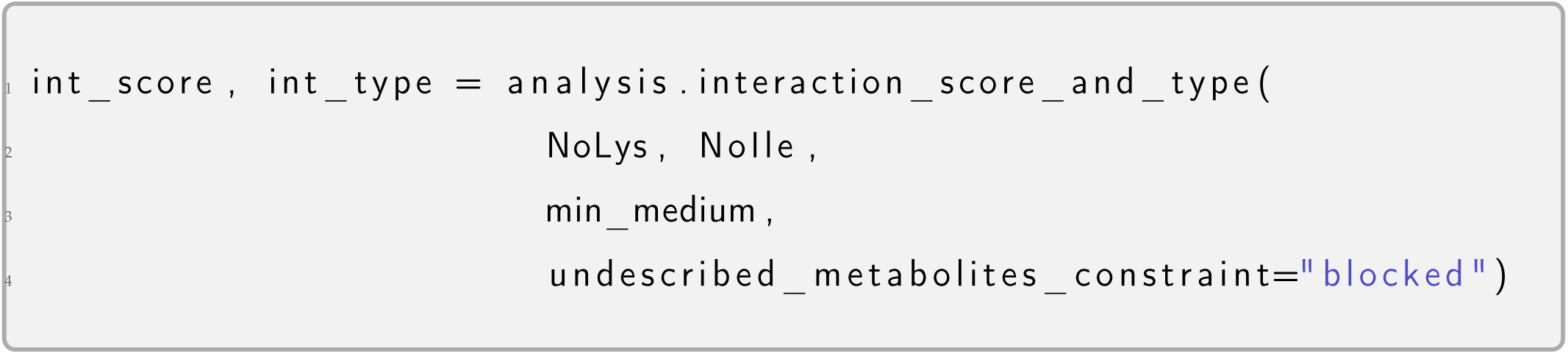

The prediction of crossfed metabolites (presented in Table **2** and Supplementary material) requires informing the identifier of the objective reaction of each model, as well as the used solver, in addition to previously required input. The crossfeeding of lysine and isoleucine was predicted using:

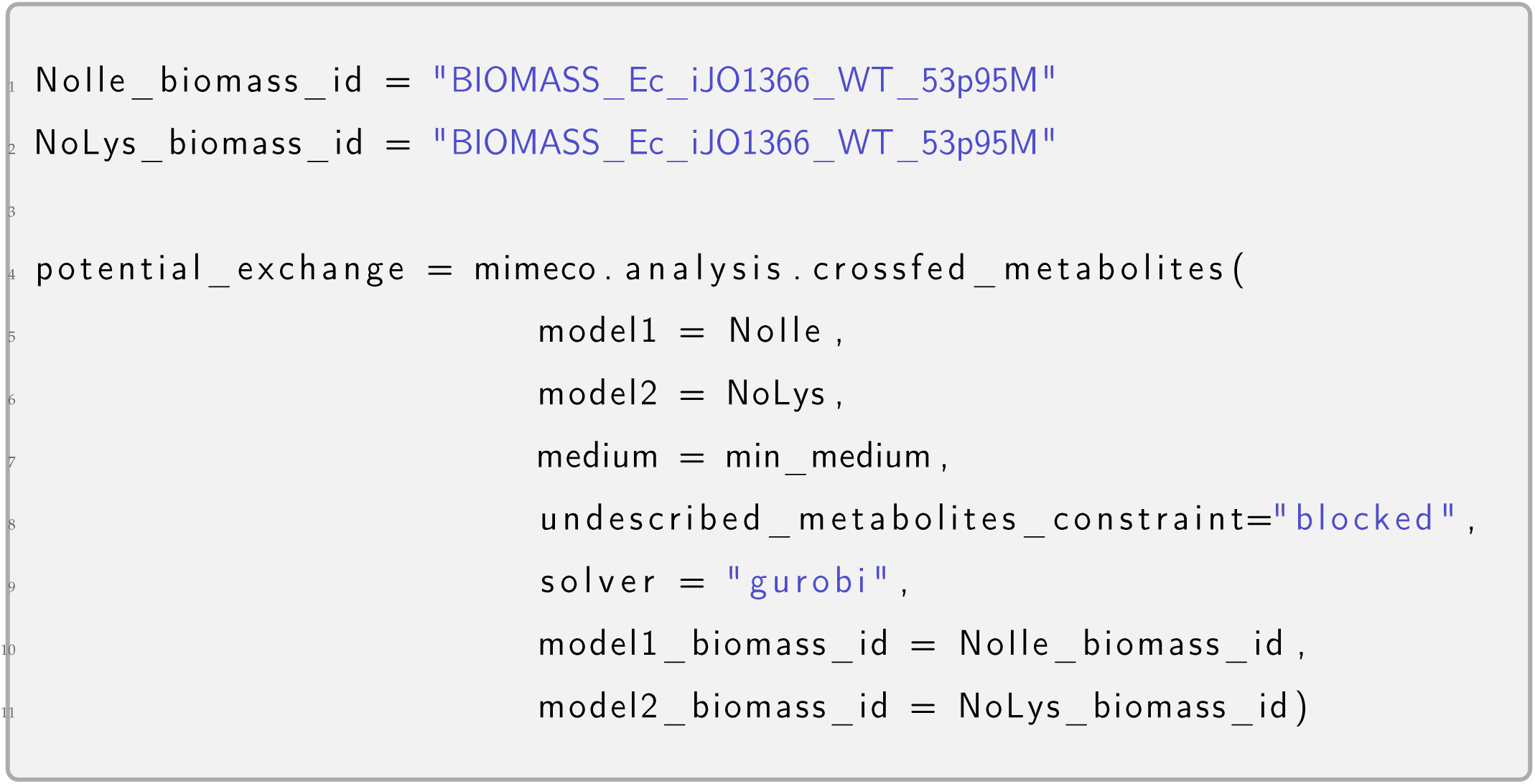

## ACKNOWLEDGMENTS

We want to thank Laura D’Hervé and Marion Morchain for using MIMEco in its development stage, giving precious feedback and allowing the improvement of the tool to fit a variety of uses. Conceptualization, A.L., D.E., and S.C.; Methodology, A.L., D.E., and S.C.; Software, A.L.; Validation, A.L.; Investigation, A.L.; Writing — original draft, A.L.; Writing — review and editing, A.L., D.E., S.C.; Visualization, A.L.; Supervision, D.E., S.C.

## DATA AVAILABILITY STATEMENT

The source code of MIMEco’s package is freely available at https://github.com/Anna-cell/mimeco Documentation with examples to install and use MIMEco is available at https://mimeco.readthedocs.io/en/latest

The scripts to reproduce the validation presented in this paper, and the script for the adaptation of the small enterocyte models are available in https://github.com/Anna-cell/mimeco/paper

---

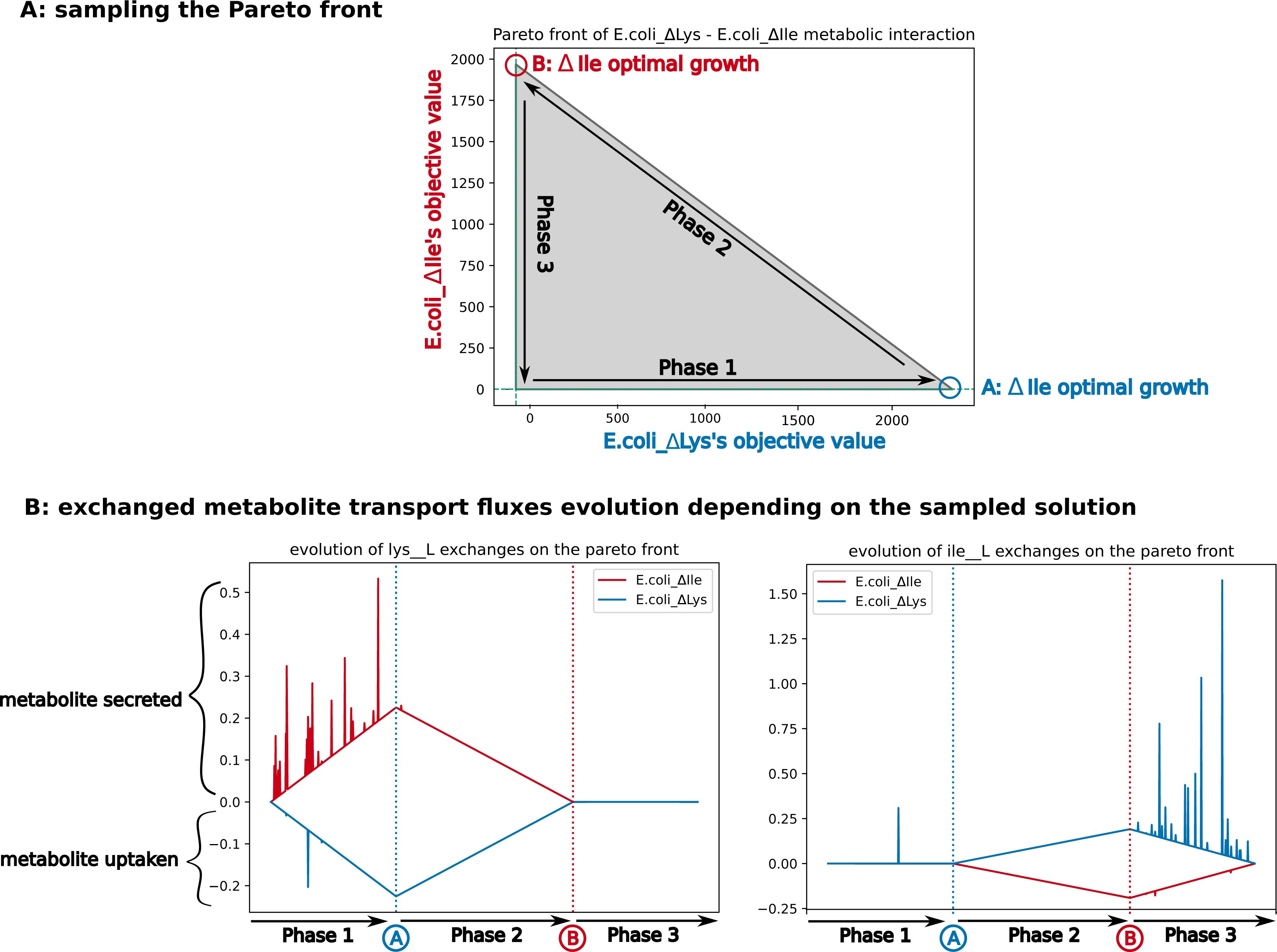

